# Inherent limitations of zero-shot mutation prediction in protein engineering

**DOI:** 10.64898/2026.06.04.730121

**Authors:** Phillip R. Woolley, Aaron L. Feller, Andrew D. Ellington, Claus O. Wilke

**Author notes:** These authors contributed equally to this work (Phillip R. Woolley, Aaron L. Feller).

## Abstract

Predicting mutation effects is critical for protein engineering. A commonly pursued strategy is zero-shot prediction with deep-learning models, whose performance is typically evaluated using aggregate benchmark scores that obscure multidimensional biological tradeoffs. Here, we evaluate sequence, structure, and hybrid foundation models across curated datasets to expose the practical limitations of zero-shot predictions. We demonstrate that model performance is highly variable and heavily influenced by experimental noise and competing phenotypic goals. Notably, predictions from competing models often correlate more strongly with one another than with experimental outcomes. While current models effectively filter deleterious mutations and enrich high-scoring variants for natural functions, they struggle to identify precise optimal mutations and consistently fail to predict mutations that drive new-to-nature functions. These models learn from evolutionary constraints that tend to oppose the extreme biophysical optimizations required in targeted engineering. Consequently, zero-shot scoring serves as a valuable initial filter but cannot substitute for task-specific prediction.

**THE BIGGER PICTURE:** Deep learning provides a powerful toolkit for predicting mutational effects in protein engineering. However, realizing the full potential of deep-learning models requires a shift in how we evaluate them. Current benchmarks report aggregate scores across many proteins, and this practice masks high variability in performance across different proteins. Furthermore, because models are trained on natural sequences, their predictions reflect evolutionary compromises rather than targeted goals. Consequently, many models struggle to predict mutations that push proteins beyond what nature requires or that introduce entirely new-to-nature functions. The next breakthrough in computational design will rely on objective-specific evaluations rather than simply scaling model architectures. Refining our metrics for success will directly enable the next generation of targeted biotechnologies.

**HIGHLIGHTS:**

- Aggregate benchmark scores mask high protein-to-protein variability.
- A single prediction score cannot decouple competing stability and activity traits.
- Model rankings enrich functional variants but miss exact optimal mutations.
- Models consistently fail to predict mutations for new-to-nature functions.

## INTRODUCTION

Predicting mutation effect across diverse proteins is a central goal in biology and medicine. The landscape of possible mutations for a single protein is vast, making exhaustive experimental characterization impractical, while computational approaches offer a scalable alternative^1–3^. Deep-learning models trained on large protein datasets have shown promise in interpreting disease-associated variants^4^ and for engineering proteins with desired properties^5,6^. Despite reported successes, the design and implementation choices of predicting mutation effect have not been standardized.

Modern methods often take advantage of a foundation model that has been trained on one or multiple sources of biological information; these may include protein sequence^7,8^, three-dimensional structure^9^, or multiple sequence alignments^10^. These models are then often evaluated using what is known as zero-shot prediction^11^, where the foundation model is used to predict the likelihood of an amino acid at each position in a protein chain. The success of a model is determined by comparing the model predictions to ground-truth data. Large benchmark datasets have been curated for this purpose, including ProteinGym^12^, FLIP2^13^, and FireProtDB^14^. In many benchmarks, performance is reported as a single aggregate score, often the average Spearman correlation between predictions and experimentally measured fitness values. These summary scores are now the primary basis for comparing models^7^, shaping both new model development and practical deployment decisions.

Even though widely used, existing benchmarks possess significant limitations. Single summary statistics compress highly variable outcomes, obscuring fluctuations in performance across different proteins and phenotypes^15^. Inherent noise in experimental measurements may impact the reliability of a benchmarking dataset^16^. Measured phenotypes frequently reflect trade-offs among competing biophysical traits; consequently, a single mutation may stabilize a fold while impairing catalytic rate or expression, producing composite effects that cannot be resolved by a single scalar prediction^17–19^. Finally, the bulk of data that ends up in benchmarks draws almost exclusively from experiments measuring natural protein function or minor variations thereof^20^. As a consequence, function-enhancing mutations and mutations that drive new-to-nature function^21,22^ are largely absent from existing benchmarks.

To address the limitations of existing benchmarking projects, here we probe specific evaluation conditions that are generally overlooked. We evaluate a curated list of foundation models trained across protein sequence and structure to explore differences in performance across specific subsets of the available data. We first evaluate standard Spearman correlation on two benchmark datasets, ProteinGym and FireProtDB. We then curate matched subsets of ProteinGym datasets^23–48^ to assess the performance of zero-shot predictions on either (i) multiple phenotypes measured for the same protein (i.e., phenotypic variability) or (ii) independent measurements of the same phenotype for the same protein (i.e., experimental noise). We further evaluate models on new-to-nature functions, where evolutionary signals are least informative^49–54^.

We find that model performance is highly variable and dataset-dependent even under standard benchmarking conditions. Our results indicate that aggregate performance masks high protein-to-protein variability, pointing to the need for evaluation frameworks designed around specific engineering goals rather than generalized performance averages. Although model rankings enrich functional variants relative to random selection, they show limited capacity to discriminate among top-tier candidates within practical screening budgets. Ultimately, these results underscore a fundamental mismatch: the balanced constraints of natural evolutionary landscapes that these models are trained on do not reliably align with the specialized objectives of targeted protein engineering.

## RESULTS

Zero-shot mutation prediction relies on the assumption that foundation models accurately learn the constraints defining a protein fitness landscape. There is a fundamental flaw in this premise that cannot be overcome by training better models or using larger training corpora: Protein fitness landscapes are multi-dimensional. Mutations that increase fitness along one dimension may decrease it along other dimensions. For example, a mutation that increases enzymatic activity often has a negative impact on protein stability^55^. A zero-shot prediction by definition assigns only a single fitness score to such a mutation, making it unable to account for the multiple factors the mutation may affect. Thus, even the best zero-shot predictions will display varying accuracy over a range of different phenotypes of interest (Fig. 1A).

**Figure 1:**
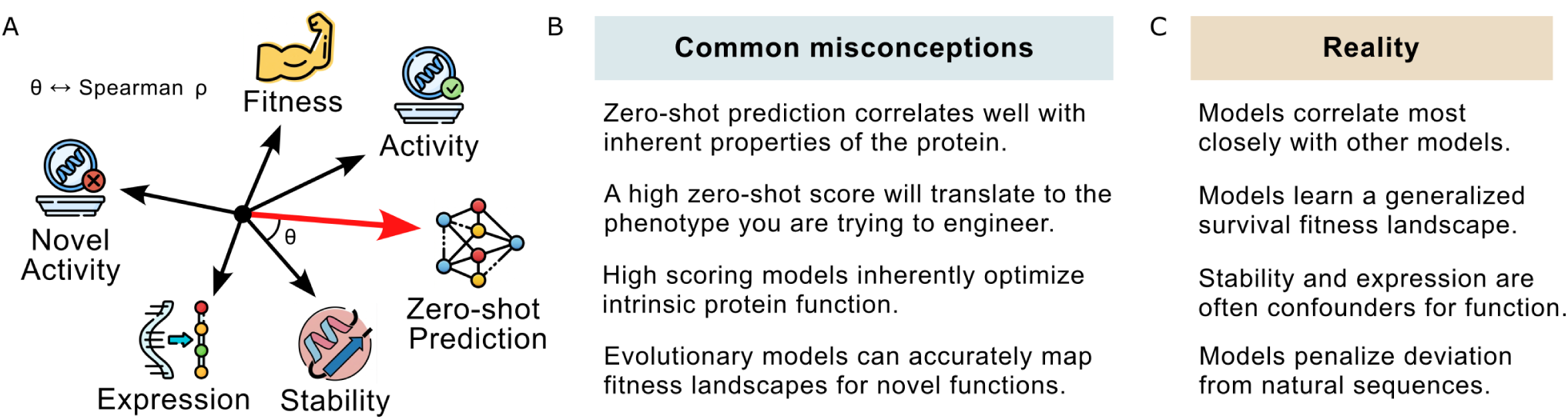
Conceptual limitations of zero-shot mutation prediction in protein engineering. (A) Zero-shot predictions cannot simultaneously capture all dimensions of a protein’s fitness landscape. A mutation that increases stability may decrease fitness and vice versa, but zero-shot prediction cannot capture these conflicting constraints. It will provide a single prediction that may or may not align with any particular phenotype of interest. In the diagram, this is indicated via the angle *θ*, which is meant to represents the Spearman correlation *ρ* between different phenotypes and/or phenotypes and zero-shot predictions. (A small angle indicates high correlation.) (B) Common misconceptions surrounding zero-shot predictions. (C) The practical reality of zero-shot modeling observed here, directly refuting the assumptions in panel B.

The existing literature on zero-shot mutation prediction largely ignores the inherent multidimensionality of protein fitness landscapes and instead operates from a basis of common misconceptions (Fig. 1B). These misconceptions are particularly problematic in the context of targeted protein engineering, where the objectives pursued may be orthogonal or detrimental to the selective pressures of organismal survival that have shaped the landscape of available data used in training foundation models. One prime example that is not captured by evolution, yet is often the goal of an engineering campaign, is improving enzyme turnover rates far beyond the need of life-sustaining processes. Any new-to-nature catalysis sought by bioengineering is absent by definition. It is realistic to expect zero-shot predictions to perform particularly poorly on such phenotypes.

Model scores reflect an aggregate history of multiple selective pressures. Beyond obvious carve-outs for targets outside the natural evolutionary landscape, single scalar predictions cannot decouple competing biophysical traits such as stability and activity. Broad benchmark averages hide these underlying conflicts. Here, we evaluate standard sequence and structure models across curated, phenotype-specific datasets. Our results outline practical limits for zero-shot prioritization and highlight critical blind spots in current benchmarking practices (Fig. 1C).

### Zero-shot model performance is highly variable across model type, dataset, and phenotypic tasks

For our analysis, we selected a panel of models representing sequence-only, hybrid, and structure-based modalities (Table 1). Rather than exhaustively evaluating all models available on public leaderboards, models were prioritized based on open-weight availability, widespread community adoption, and computational tractability for scoring large-scale mutational libraries. We included ProteinMPNN and SolubleMPNN, even though they were originally designed for backbone-conditioned sequence generation, because conditional log-likelihood ratios from inverse-folding models are routinely deployed as zero-shot mutational effect predictors in the literature^56^. All models were run in a zero-shot setting without task-specific fine-tuning.

**Table 1:** Models evaluated for zero-shot performance in this study. The models include pure sequence-based models, mixed models that take advantage of both sequence and structure data, and pure structure-based models that do not consider sequence.

| Model Category<sup>1</sup> | Model Name | Parameters<sup>2</sup> | Citation<sup>3</sup> |
| --- | --- | --- | --- |
| Sequence-based | AMPLIFY | 350M | 57 |
|  | ESM-2 | 650M, 3B | 8 |
|  | ESMC | 300M, 600M | 58 |
| Mixed sequence/structure | ESM-3 Open | 1.4B | 59 |
|  | ISM | 650M | 60 |
|  | ISMC | 300M, 600M | 60 |
|  | SaProt | 650M | 61 |
| Structure-based | ProstT5 | 3B | 62 |
|  | ProteinMPNN | 1.7M | 9 |
|  | SolubleMPNN | 1.7M | 63 |

<sup>1</sup> Input data type of the model.
<sup>2</sup> Number of parameters of the model. If multiple numbers are listed all listed variants were evaluated.
<sup>3</sup> Publication originally describing each model.

To determine how input modality and target phenotype influence zero-shot performance, we compared model rankings (Spearman *ρ*) on two benchmark datasets: The FireProtDB thermostability benchmark^14^ and the ProteinGym fitness benchmark^12^. When sorting models by their median performance (Spearman *ρ*), we found that the models that performed best on stability performed poorly on fitness, and the top models for fitness also had decreased performance on stability (Fig. 2A). Moreover, when categorizing models into sequence-only, structure-only, or hybrid modalities, we observed a slight advantage for hybrid models in fitness ranking but a mixed pattern for structure-conditioned models in thermostability ranking (Fig. 2A). Overall, we found that all models performed poorly on some datasets and excellent on others (*ρ* ranging from less than zero to near one), while the median performance differed only moderately among models (Fig. 2A) (median *ρ* ranging between 0.3 and 0.5 for all models and both benchmarks).

**Figure 2:**
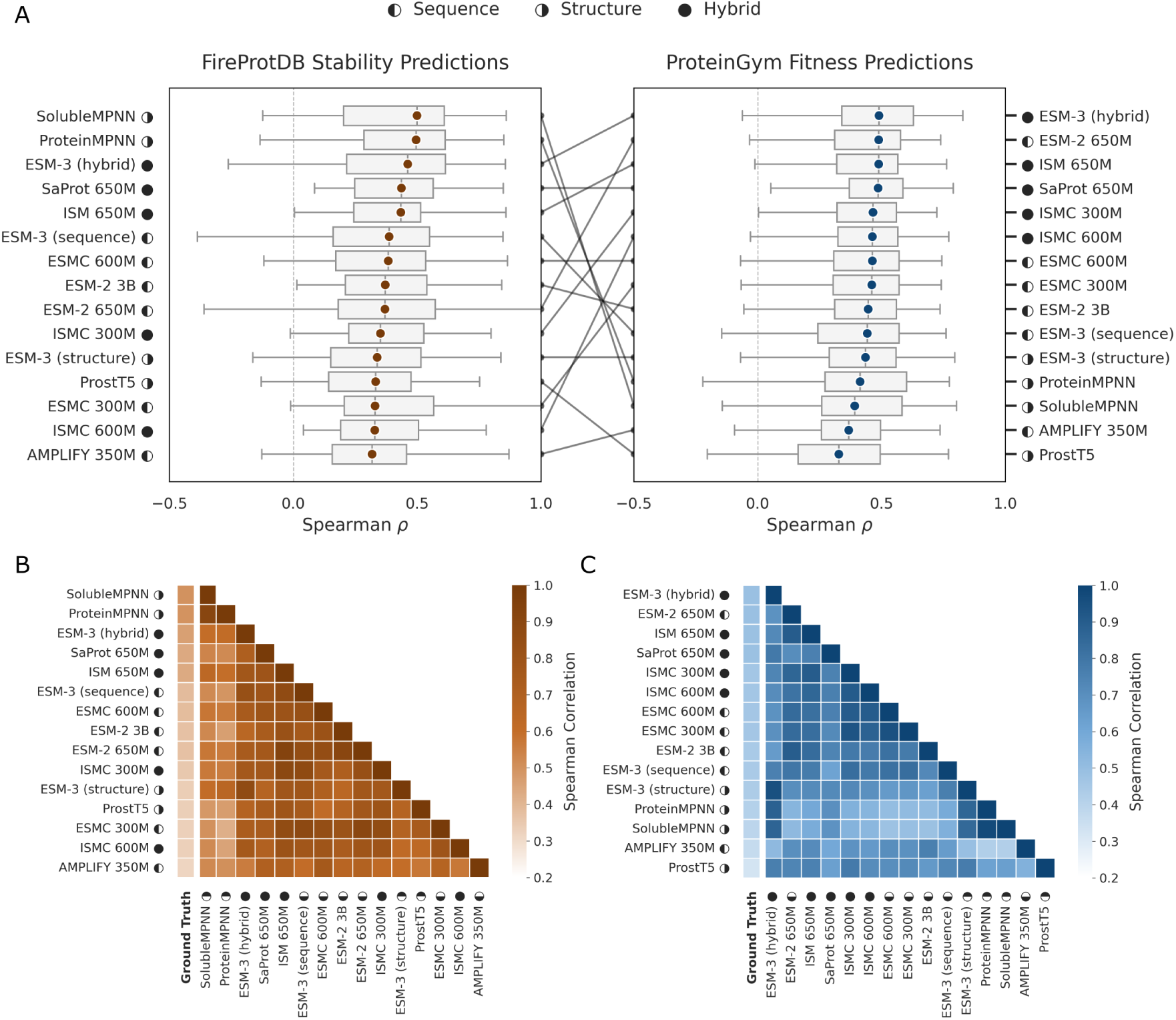
Zero-shot model performance is weakly predicted by input modality and highly variable across datasets and phenotypic tasks. (A) Zero-shot model performance (Spearman *ρ*) on the FireProtDB thermostability benchmark (left) and the ProteinGym Fitness benchmark (right), ordered by median *ρ*. Each boxplot shows the median, interquartile range, and 1.5*×*IQR whiskers over all test proteins, with colored circles representing the median. The vertical dashed line indicates random performance (*ρ* = 0). (B) Average ranked correlation (*ρ*) between each model and the ground truth and pairwise between all models for all proteins in FireprotDB with *>* 20 mutations. (C) Average ranked correlation (*ρ*) between each model and the ground truth and pairwise between all models for all proteins in ProteinGym.

To determine whether these performance differences were consistent at the dataset level, we compared model performance on individual datasets. Despite achieving comparable aggregate scores, models displayed significant disagreement on individual datasets within the same phenotype class (Fig. S1). We quantified these performance differences by calculating Spearman correlations between the *ρ* distributions of each model across individual datasets in FireProtDB and ProteinGym. The correlation coefficients ranged from *−*0.61 to 1.0 in FireProtDB (Fig. 2B) and from *−*0.36 to 0.83 in ProteinGym (Fig. 2C). These findings indicate that zero-shot performance is highly dataset-dependent, even among top-performing models, and that aggregate benchmark scores obscure meaningful divergence in model behavior. Consequently, relying solely on aggregate performance is insufficient for selecting the optimal model for a specific protein and task.

### Mutational effect measurements vary between laboratories and phenotypic objectives

To isolate potential sources of variation in model performance, we investigated inter-experimenter and inter-phenotype effects. We established two controlled comparisons using subsets of the ProteinGym benchmark (Table 2): (i) independent laboratories assaying a single mutation set for the same objective and (ii) a single research group evaluating the same protein across multiple phenotypic objectives.

**Table 2:**
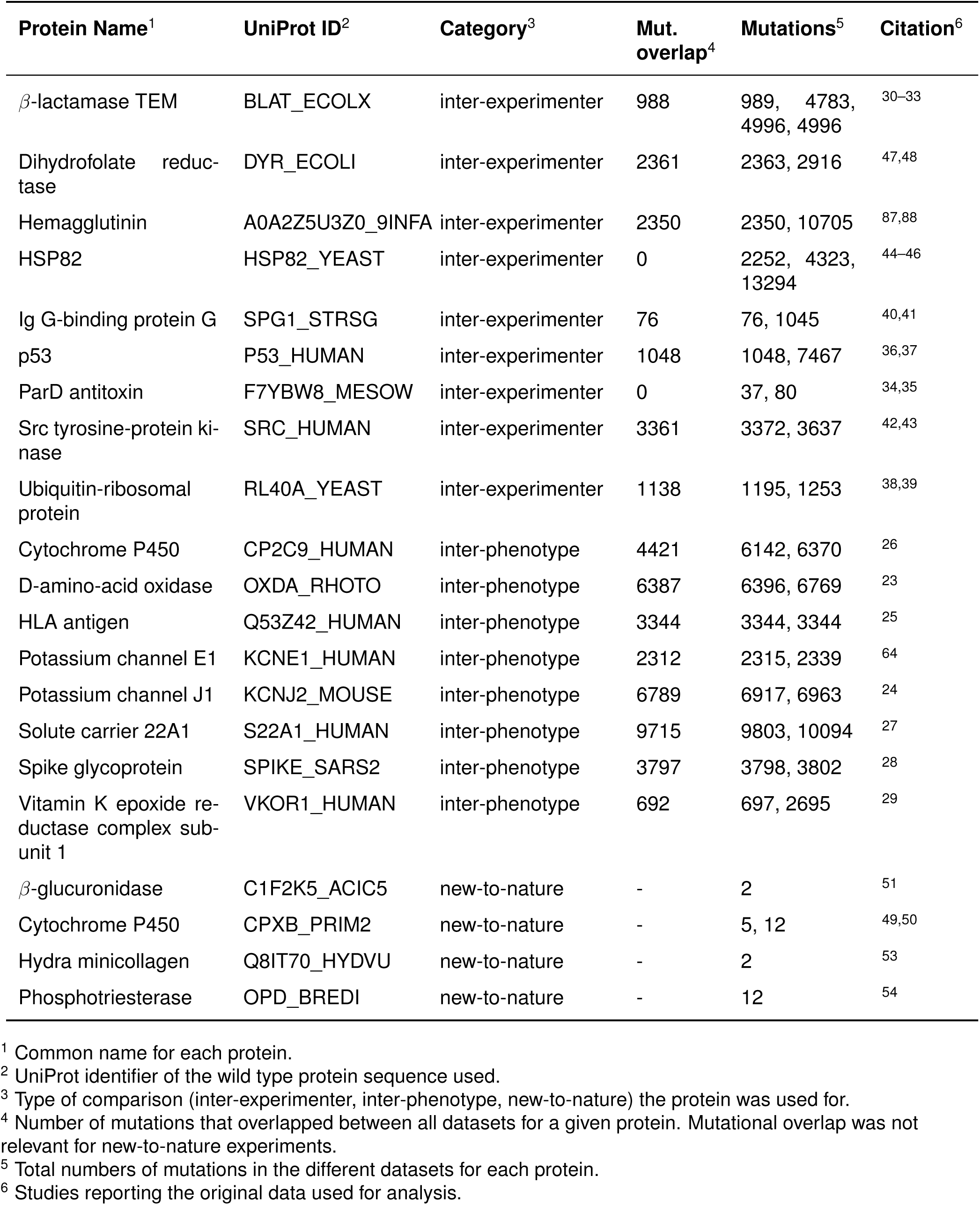
Datasets used for inter-experimenter, inter-phenotype, and new-to-nature comparisons.

Comparisons between independent laboratories assaying the same phenotype revealed moderate to strong correlations among the fitness measurements (*ρ* = 0.55–0.9; *n* = 76–3361 overlapping mutations per dataset). However, the degree of agreement and mutational overlap between datasets varied considerably across proteins (Figs. 3A–C and S2, Table 2). This observation confirmed that experimental noise and methodological differences introduce a moderate degree of divergence in measured mutational effects.

**Figure 3:**
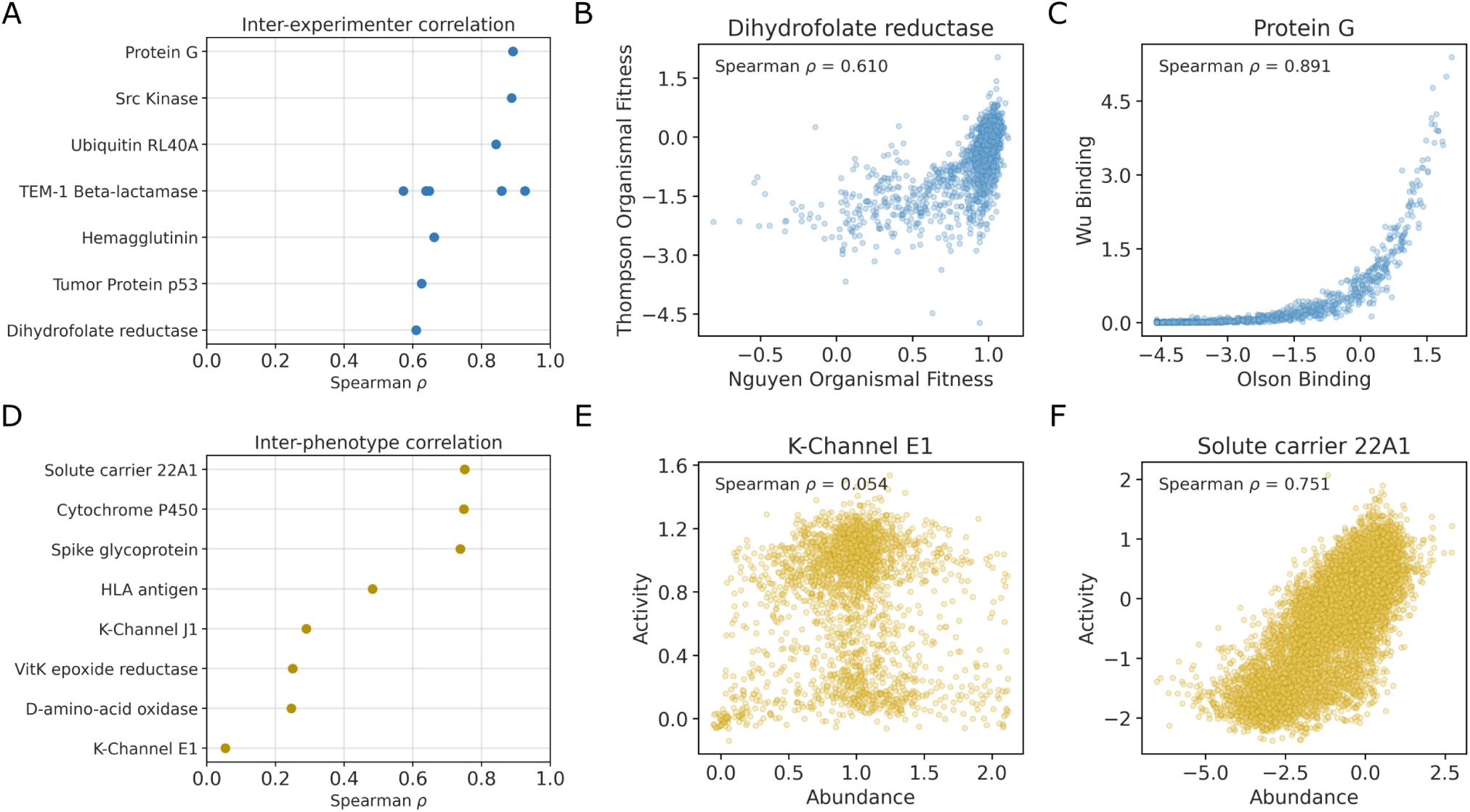
In ProteinGym datasets, inter-experimenter agreement is moderate, while inter-phenotype correlations are highly variable. The top three panels show inter-experimenter comparisons, in which two independent groups assayed the same protein for the same phenotypic objective. The bottom three panels show inter-phenotype comparisons, in which a single group assayed the same protein for two distinct phenotypes; in every such case within ProteinGym, one of the two phenotypes is protein abundance. (A) Spearman *ρ* for all inter-experimenter comparisons available in ProteinGym. (B) Scatter plot of the inter-experimenter pair with the lowest Spearman *ρ* (on average), dihydrofolate reductase^47,48^ (C) Scatter plot of the inter-experimenter pair with the highest Spearman *ρ* (on average), protein G^40,41^. (D) Spearman *ρ* for all inter-phenotype comparisons available in ProteinGym. (E) Scatter plot of the inter-phenotype pair with the lowest Spearman *ρ*, potassium channel E1^64^. (F) Scatter plot of the inter-phenotype pair with the highest Spearman *ρ*, solute carrier 22A1^27^.

Conversely, inter-phenotype comparisons exhibited a much broader range of correlations (*ρ* = 0.05–0.8; *n* = 692–9715 overlapping mutations per dataset) (Figs. 3D–F and S3, Table 2). Notably, in every ProteinGym study measuring multiple objectives for a single mutation set, one reported phenotype was protein abundance. The other measured phenotypes typically reflected aggregate protein performance across the cellular system, integrating contributions from expression, folding, trafficking, and activity^23–29^. This observation points to an important caveat in comparing multiple phenotypic measurements for the same protein: mutations increasing protein abundance may appear to improve functional performance without altering intrinsic catalytic rate, substrate affinity, or specificity. Disentangling these variables experimentally would require precise measurements that remain difficult to obtain at scale.

The relationship between abundance and functional scores across multi-phenotype studies (Fig. 3D–F) is further complicated by limitations in assay resolution. For instance, potassium channel E1 sets the lower bound for these correlations, displaying no correlation at all between activity and abundance (Spearman *ρ* = 0.054, Fig. 3E). The lack of correlation can be explained by the functional assay used, which is effectively binary^64^, where any active variants enhance a toxic background that results in cell death. This binary readout compresses the resolution of the functional phenotype, limiting meaningful separation among active mutations. Conversely, in cases where abundance and functional scores are highly correlated, apparent gains may largely reflect improvements in biophysical properties of the expressed protein, such as solubility and stability, rather than an improvement in the native function of the protein.

### Inter-experimenter and inter-phenotype variance propagates into model performance assessments

Model rankings in benchmarking studies are highly sensitive to the choice of experimental dataset. For ProteinGym datasets sharing the same protein and reported phenotype, the impact of dataset differences on model performance ranged from negligible to substantial (Fig. 4A). This effect was inconsistent across proteins and models: For some proteins, the difference in inter-experimenter performance for most models was minimal, while for other proteins (e.g., Protein G) the difference in *ρ* exceeded 0.6 for some models. Proteins with the largest inter-experimenter effects tended to have low mutational overlap between datasets, suggesting that model performance was strongly influenced by which regions of the protein were sampled. The mean absolute difference in model performance for inter-experimenter and inter-phenotype comparisons revealed that lower mean absolute difference did not imply stronger aggregate performance (Fig. 4B). In particular, the model AMPLIFY 350M had the lowest inter-experimenter mean absolute difference (Fig. 4B) but performed poorly overall (Fig. 2B).

**Figure 4:**
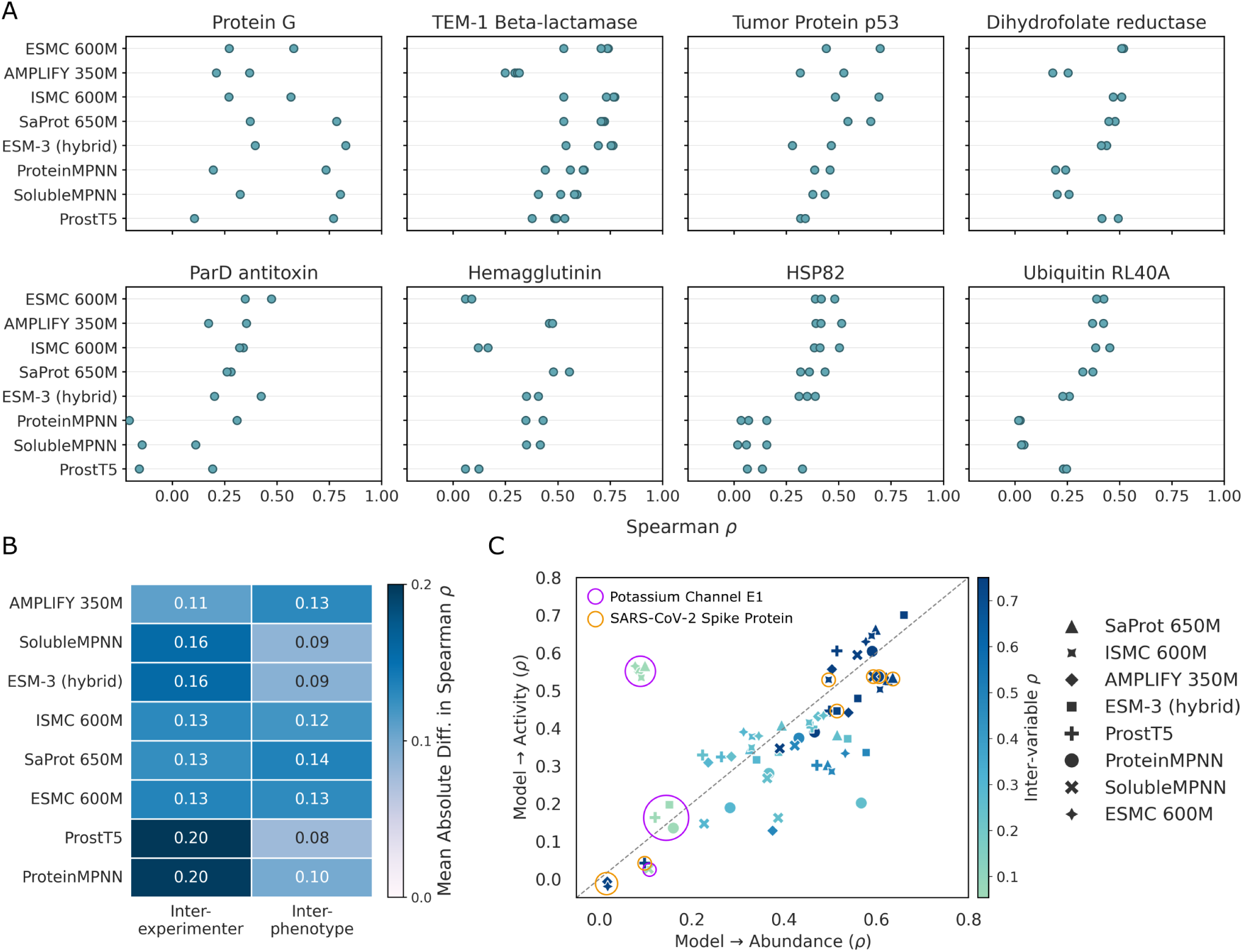
Variability in experimental dataset and phenotype leads to variability in model performance assessments. (A) Spearman *ρ* for eight proteins with multiple experiments for the same measured phenotype. Each point represents one *ρ* value for one specific experiment/model combination. (B) Mean absolute difference between the largest and the smallest Spearman *ρ* across all inter-experimenter and all inter-phenotype comparisons, for each model. (C) Spearman *ρ* measuring model performance in predicting functional activity (*y*-axis) versus protein abundance (*x*-axis) for datasets in the multiple-phenotype group. Each point represents a single model’s *ρ* on each phenotype. Point color indicates the Spearman *ρ* between the abundance and activity phenotype scores, and point shape indicates the model. Purple and orange circles highlight model performance on Potassium Channel E1 and SARS-CoV-2 Spike Protein, respectively.

For inter-phenotype comparisons, model performance across the two phenotypes generally tracked the degree to which the phenotypes themselves correlated (Fig. 4C). When abundance and functional scores agreed, models performed well on both; when the phenotypes disagreed, models performed poorly on both. However, there was a modest bias for models to perform better when predicting abundance than when predicting activity, as can be seen from the fact that there are more points below the diagonal than above in Fig. 4C.

We note two proteins that deviated from this overarching trend, Potassium Channel E1 and SARS-CoV-2 Spike Protein (Fig. 4C). Potassium Channel E1 displayed model scores split into two tight clusters (either performing well on the functional phenotype or poorly on both). In other words, for this protein, some models could predict activity well and others couldn’t, while no models could predict abundance well. For SARS-CoV-2 Spike Protein, models either performed well on both phenotypes or poorly on both. The likely cause of poor model performance in this case was absence of this viral protein from the models’ training corpora^65,66^. Excluding these two anomalies, the overarching pattern we found suggests that phenotype correlation can serve as a rough guide to expected model behavior.

### Models enrich high-scoring activity variants but do not reliably recover the optimum

To evaluate the utility of zero-shot methods for protein engineering campaigns, we identified mutations that were highly scored by zero-shot models and asked how many of them actually resulted in increased experimentally measured activity. For this analysis, we selected every ProteinGym dataset annotated as an Activity assay and ranked all assayed single mutations by model log-odds. As an example to illustrate our approach and the relationship between model scores and experimental outcomes, we have visualized the S22A1 activity landscape (Fig. 5A). We split mutations into categories based on measured activity, corresponding to worse than wild type, beneficial, top 10%, and top 1% of measured activity. We then evaluate the ability of the model ranking to recover targets from these categories across increasing screening budgets. The idea here is that in cases where experimental testing of mutations is costly, the budget available for screening may be limited, and we may only be able to subject a limited number of mutations to experimental testing, such as 32 or 64. The question then becomes whether we have a reasonable chance of finding an improved mutation among the tested ones, and whether zero-shot predictions help increasing the odds of discovering improved mutations. Because we performed this analysis retrospectively on known, already experimentally measured single mutants, it is somewhat limited, in particular for datasets that did not exhaustively test all possible single-point mutations.

**Figure 5:**
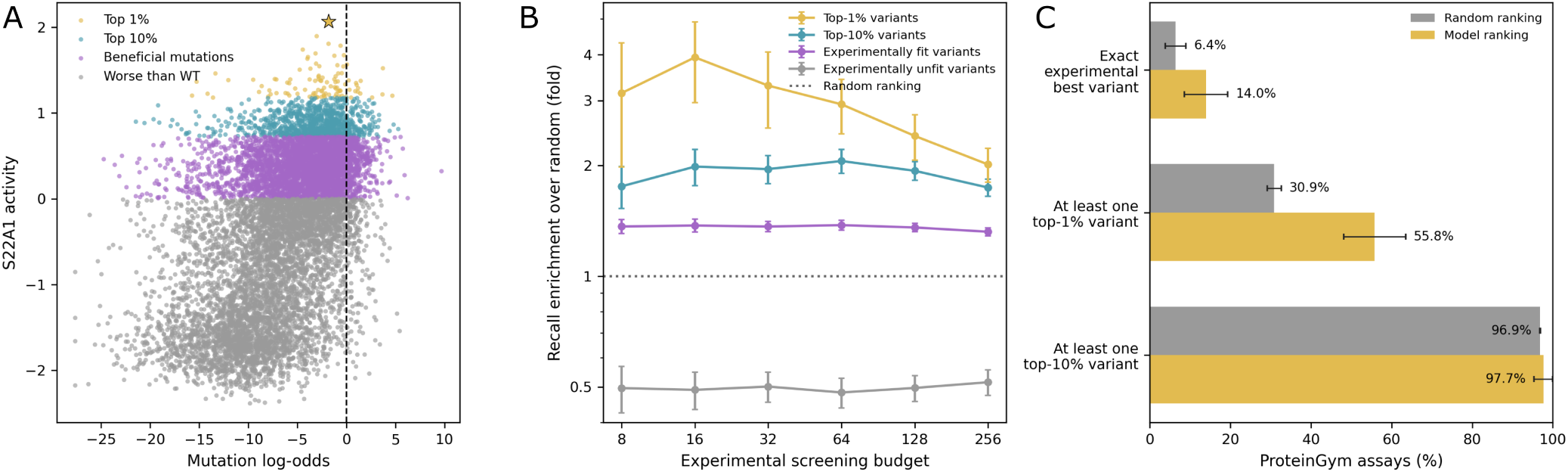
ESM-3 (hybrid) ranking enriches experimentally fit and high-scoring variants in ProteinGym Activity assays. (A) Experimental activity versus mutation log-odds from ESM-3 (hybrid) for solute carrier 22A1 (S22A1)^27^. Grey points are worse than wild type, purple points are remaining beneficial mutations, blue points are in the top 10% of activity values, and gold points are in the top 1%; the star marks the experimentally best variant. The vertical dashed line indicates zero log-odds. (B) Selection enrichment, defined as the ratio of model-ranked recall to the expected recall from random screening, across six screening budgets. Error bars are standard errors across assays. (C) At a fixed budget of 32 mutations, the fraction of Activity assays in which random or model ranking recovers the exact experimental best variant, at least one top-1% variant, or at least one top-10% variant. Error bars are standard errors across all fitness assays.

We evaluated three models with this screening protocol; the ESM-3 as the top hybrid model for predicting fitness (Fig. 5), ProteinMPNN as the top structure model for predicting stability (Fig. S4), and ESMC-600M as the top sequence-based model (Fig. S5). All three models enriched the target groups above random screening over most displayed budgets (Figs. 5B, S4B, S5B). At a budget of 32 candidates, both random selection and ESM-3 (hybrid) recovered at least one top-10% activity variant in nearly all of the assays (>95%), and at least one top-1% variant in 55.8% of assays, compared with 30.9% at random (Fig. 5C). However, recovery of the exact experimental best variant was low (14.0% for ESM-3 versus 6.4% at random). Thus, zero-shot scores provide useful, but heterogeneous, enrichment for high-scoring activity variants in a constrained screening setting.

### Protein models are poorly suited for zero-shot prediction of new-to-nature functions

New-to-nature tasks constitute a major subset of protein engineering challenges, encompassing cases where a novel function is accessible by a small number of mutations to a naturally existing protein. Because protein models are trained on naturally evolved sequences, their learned probability distributions reflect evolutionary fitness, capturing native function and general biophysical properties such as solubility and stability. Mutations that redirect a protein toward a new-to-nature function fall outside the evolutionary context these models have learned, and we expect model performance on such mutations to be poor.

To test this hypothesis, we collected examples from the protein-engineering literature where a minimal set of mutations was sufficient to redirect a protein toward a novel functional objective (Table 2)^49–51,53,54^. We evaluated three models for this analysis, the top performing model for each modality: ESM-3 (hybrid) (Fig. 6), ESMC 600M (Fig. S6) and ProteinMPNN (Fig. S7). Across all five datasets combined (*N* = 33 mutations, *z*-score normalized), all three models showed no positive correlation with measured phenotype (Spearman *ρ ≈ −*0.3; Figs. 6A, S6A, S7A).

**Figure 6:**
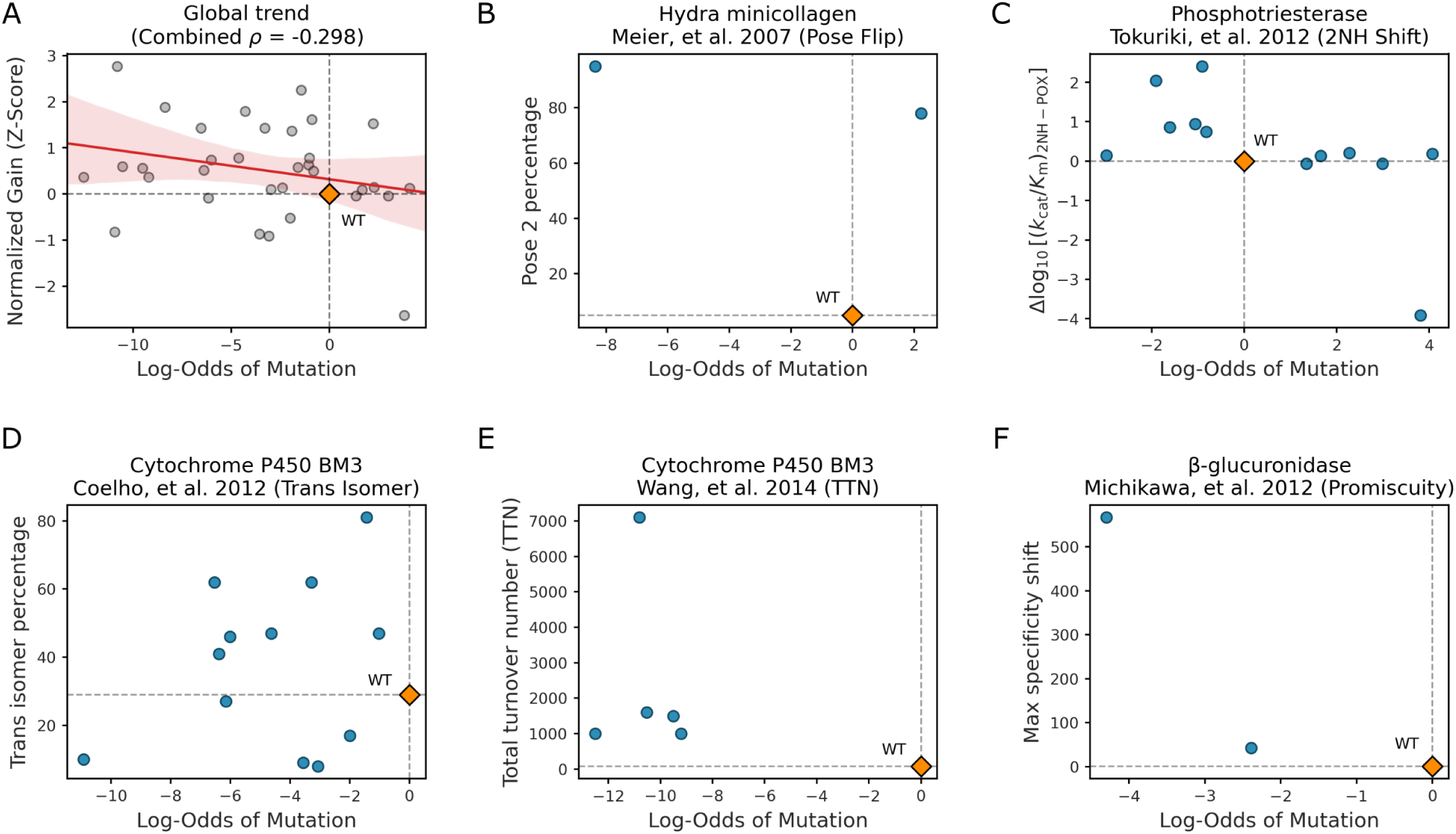
ESM-3 (hybrid) performs poorly on new-to-nature function mutations. (A) Pooled results from five examples of new-to-nature engineering campaigns. Each point represents a single mutation. Normalized gain indicates the fitness increase (or decrease) of a mutation relative to wild type, represented as a *z*-score to make results comparable across the five experimental systems. Log-odds is the model prediction for each mutation. Overall, there is a slight negative correlation between normalized gain and log-odds, i.e., mutations predicted to be better perform on average worse. (B) Hydra minicollagen engineering to favor an alternative conformational state^53^. (C) Phosphotriesterase engineering to favor an alternative substrate^54^. (D) Cytochrome P450 engineering to optimize trans isoform percentage^49^. (E) Cytochrome P450 engineering to optimize total turnover number (TTN)^50^. (F) *β*-glucuronidase engineering for promiscuity toward alternative substrates^51^. Similar results for models ESMC-600M and ProteinMPNN are shown in Figs. S6 and S7, respectively.

When reviewing model predictions for individual datasets, we identified consistent model failure across all tasks. In one case, a D-hydantoinase microcollagen antigen with a conserved disulfide-governed domain adopted distinct conformational isoforms depending on the residues surrounding the disulfides^53^. Starting from a wild-type sequence predominantly in state 1, a single point mutation experimentally shifted the equilibrium toward state 2, and a second mutation drove the protein to adopt state 2 almost exclusively. Despite this clear trajectory, ESM-3 (hybrid) favored only the first mutation (Fig. 6B), while ESMC 600M (Fig. S6B) and ProteinMPNN (Fig. S7B) incorrectly disfavored both. In a second example, a phosphotriesterase engineering campaign progressively shifted substrate specificity from Paraoxon to 2-naphthyl hexanoate across a series of point mutations^54^. Using the previous mutation round as the sequence background for each successive mutation, model predictions remained poor (Figs. 6C, S6C, S7C). Two datasets were from a cytochrome with a catalytic cis:trans product ratio that was tuned through mutation. Model performance was poor, whether the objective was enriching trans-product formation^49^ (Figs. 6D, S6D, S7D) or optimizing turnover number^50^ (Figs. 6E, S6E, S7E). Finally, for mutations in *β*-glucuronidase identified as broadening substrate promiscuity^51^, ESM-3 (hybrid) and ESMC 600M both rejected both mutations over wild-type (Figs. 6F, S6F), while ProteinMPNN rejected one and barely favored the other (Fig. S7F).

For the phosphotriesterase engineering campaign, where mutations were introduced sequentially, we also modeled all mutations individually in the wild-type background, to assess whether mutation context affected performance. We found nearly identical results to using the previous mutation round as the sequence background (Fig. S8). Taken together, these results demonstrate that the predictive ability of current protein models does not reliably extend to new-to-nature contexts, a finding consistent across all three models tested.

## DISCUSSION

Deep-learning models are now standard tools for predicting mutational effects, routinely benchmarked using average scores across curated datasets^12,14^. However, summary metrics obscure critical, context-dependent performance variation; notably, the median scores of competing models vary less than the performance of any single model across different datasets. Consequently, median metrics offer little predictive power for how a model will perform on a specific protein. More elemental, experimental measurements vary significantly across independent laboratories even for identical mutations^67^. Furthermore, aggregate scores are easily skewed by assay-based confounding effects; high-throughput assays frequently fail to normalize for protein abundance^68–70^, causing models to appear successful at predicting function when they are merely predicting stability. Even without these confounding effects, different phenotypes measured for the same mutations may be non-correlative.

Failure modes are abundant. When mutations exert conflicting effects across distinct biophysical traits, one zero-shot score cannot simultaneously track multiple objectives. Because these scores reflect a balance of traits optimized for natural survival, they inherently struggle when engineering goals require pushing a single biophysical property to an extreme. This limitation is particularly evident when evaluating alternative conformational states: because state transitions are mutually exclusive, a scalar score cannot simultaneously favor both conformations. In such cases, a low correlation does not necessarily indicate a defective model, but rather illustrates the intrinsic limit of evaluating multi-state equilibria with an unconditioned score. Further, we have found that benchmarks are easily skewed by specific outliers, such as the binary read-out of Potassium Channel E1^64^, or critical gaps in training data exposed by viral proteins like the SARS-CoV-2 spike^65,66^. The alignment between benchmark evaluations and model development warrants careful scrutiny. Models that are validated against or optimized on benchmark datasets during development risk circularity, inflating benchmark scores over genuine performance in an engineering campaign, which may ultimately reduce trust in certain models and impact model adoption.

To ensure our findings reflect the broader landscape, we have evaluated a panel of widely adopted architectures across three primary modalities: sequence-only (AMPLIFY^57^, ESM family^8,58^), structure-only (ProteinMPNN^9,63^, ProstT5^62^), and hybrid or distilled models (ESM-3^59^, ISM^60^). While ProteinMPNN was designed for inverse folding, conditional log-likelihoods from such models are routinely used as zero-shot predictors^56^. We have observed some notable differences across modalities. Purely structural MPNN models perform best on stability but worst on function, whereas structure-conditioned language models like ESM-3 perform well on both. Overall, it appears that a model incorporating protein structure is advantageous for predicting stability^71^, though we caution against over-interpreting aggregate benchmark rankings.

The convergence of median performance scores across diverse architectures points to a fundamental boundary in zero-shot prediction. We have found that model predictions correlate more strongly with one another than with experimental ground truth, a phenomenon that we suggest stems from their shared training corpora of naturally occurring proteins^2,7^. If correct, all foundation models would be operating within the constraints of evolutionary history, learning to balance the biophysical properties required for survival. Targeted engineering, however, often requires overriding these natural balances. Because mutations that dramatically enhance a single functional property often incur severe fitness costs in a natural context^17–19^, they are selected against in the wild. As a result, highly engineered variants occupy sequence space absent from the training data, causing zero-shot models to systematically score them poorly.

This limitation of zero-shot models is most pronounced when targeting new-to-nature functions. To assess zero-shot performance in this regime, we have curated several datasets from the directed-evolution literature where a protein acts on a non-natural substrate or performs novel chemistry. We have deliberately excluded thermostability or solubility traits, which are under constant natural selection. In these non-natural contexts, zero-shot models consistently fail to predict activity-enhancing mutations. Our findings are consistent with recent evidence that mutations prioritized by protein foundation models perform worse at establishing novel functions than variants selected from simple site-level variation in sequence alignments^72^. We conclude that for novel functional engineering, zero-shot models are better deployed downstream to recover baseline properties like stability or solubility after a functional variant has been identified^73–75^.

Despite these limitations, zero-shot models still hold some practical value as coarse filters. On activity readouts, model rankings preferentially enrich experimentally fit variants over random screening within finite candidate sets. However, exact recovery of top-tier mutations (e.g., the top 1%) remains inconsistent. Overcoming this barrier requires moving beyond generic evolutionary priors. When screening data is available, supervised fine-tuning and few-shot paradigms, such as those evaluated in FLIP2^13^ or via task-specific adaptation^76–78^, can anchor learned representations directly to target phenotypes. Alternatively, classical biophysical functions, such as FoldX^79^ or PRIMO^80^, alongside other physics-based methods^81–83^, can evaluate thermodynamic stability independent of evolutionary history. Moving forward, engineering pipelines would benefit from external alignment diagnostics^65,84^ and uncertainty-aware metrics^85^ to guide model selection, similar to established practices in protein structure prediction^8,86^.

Ultimately, the field of protein engineering currently lacks the rigorous, multi-phenotype experimental datasets needed to evaluate models in underrepresented contexts. Paired multi-replicate measurements from independent laboratories, deep mutational scanning datasets spanning multiple phenotypes for the same mutations, and annotated directed evolution trajectories with characterized intermediates remain scarce. Because the limitations of zero-shot prediction are rooted in the evolutionary training data itself rather than parameter counts, further investment in model scaling or novel architectures will yield diminishing returns. Moving forward, coordinated efforts to generate and share targeted experimental data will do far more to guide protein engineering than simply building larger models.

## METHODS

### Dataset selection and curation

FireProtDB (v2.0)^14^ was used as a source for thermostability data. The database provides Δ*T_m_* (the change in melting temperature) and Δ*G* (the change in folding free energy) values for single amino-acid substitutions, reported on a per-dataset basis as either one metric or the other. Where Δ*G* was available for a given dataset, it was preferred over Δ*T_m_* and multiplied by −1 so that larger values indicate greater stability; otherwise, Δ*T_m_* was used directly. Only single amino-acid substitutions were retained for analysis. FireProtDB UniProt^89^ identifiers were used to retrieve sequences and structures. In total, we collected stability data for all datasets containing at least 20 mutations, leading to a total of 4824 mutations spanning 39 distinct proteins (Supplemental File S1). We removed proteins that were unable to be encoded with all models we chose due to length/size limitations.

ProteinGym (v1.3)^12^ was used as the source of deep-mutational scanning (DMS) datasets for fitness benchmarking (Fig. 2) and for inter-experimenter and inter-phenotype analyses. We used the DMS score provided in the ProteinGym substitution tables and retained only single mutants (mutation strings without a colon, “:”). ProteinGym was used strictly as a dataset source; no ProteinGym model scores were used in any analysis. In total, we collected DMS scores for 2.47 million mutations spanning 217 distinct DMS experiments (Supplemental File S2).

We manually reviewed all the ProteinGym DMS datasets for cases in which either independent laboratories had assayed the same protein and phenotype (inter-experimenter comparisons) or in which a single research group had assayed the same protein for multiple phenotypes (inter-phenotype comparisons). We found nine cases that allowed for inter-experimenter comparisons and eight that allowed for inter-phenotype comparisons (Table 2).

For both inter-experimenter and inter-phenotype comparisons, we found that there was often a discrepancy in the exact mutations that were evaluated by different groups or for different phenotypes, and therefore we identified the mutational overlap, defined as the intersection of single-point mutations across the relevant datasets. Mutational overlap ranged from zero to 9715 across the various datasets (Table 2). We excluded the two groups of datasets with zero mutational overlap from inter-experimenter analyses in Fig. 3 but retained them in Fig. 4.

New-to-nature datasets were curated from directed-evolution studies with small sets of single-point mutations and quantitative phenotype readouts. To identify candidate papers, we queried LLM chatbots and search engines using terms such as “de-novo protein design,” “directed evolution campaigns,” and “new-to-nature functionality,” and mined review articles for references to relevant primary papers. We evaluated each candidate paper for whether it reported the effects of single point mutations on novel protein functionality, such as performing novel chemistries or acting on novel substrates. Studies focusing on general productivity-enhancing traits such as thermostability or solubility were excluded. Similarly, studies focused on swapping protein domains or otherwise making large-scale modifications, on *de-novo* design of entire proteins, or on any other modifications other than individual point mutations were also excluded. The vast majority of papers we evaluated did not meet these criteria. Deep mutational scanning studies, while comprehensive in mutational coverage, tended to characterize mutations only with respect to native protein function rather than novel tasks. On the flip side, many protein design and protein engineering studies did not report single point mutation effects.

Our final set of new-to-nature datasets include Hydra minicollagen conformational switching (Fig. 2 from^53^), phosphotriesterase substrate specificity shifts (Tables 2–3 from^54^), cytochrome P450 trans-isomer enrichment and turnover (Table 1 from^49^, Table 1 from^50^), and *β*-glucuronidase promiscuity shifts (Table 3 from^51^). These datasets are summarized in Table 2. The small number of new-to-nature datasets in our analysis reflects the limited availability of suitable data in the literature.

### Structure preparation and 3Di tokenization

For all proteins obtained from FireProtDB and ProteinGym, we required structures to run the structure-based models SaProt^61^, ProstT5^62^, and ESM-3^59^. For proteins from FireProtDB, we obtained structures from the AlphaFold Database (AF-DB)^90^ using the provided UniProt identifiers. For proteins from ProteinGym, we used the structures provided by ProteinGym where available. One protein from ProteinGym (ANCSZ_Hobbs_2022) and all new-to-nature proteins were modeled with Boltz-2^91^ using default settings and the MMseqs2 MSA server.

Foldseek 3Di tokens^92^ were generated from PDB structures using Foldseek tools createdb, lndb, and convert2fasta to create a foldseek.fasta file containing the 3Di tokens. These 3Di sequences were paired with amino-acid sequences for SaProt and used as direct input for ProstT5. For ESM-3, PDBs were parsed into structure tokens using the ESM-3 structure encoder.

### Model selection and zero-shot mutation prediction

We selected a panel of models representing different input modalities and model architectures (Table 1). Our set of model choices was not meant to be as exhaustive as possible but rather to represent common model architectures and widely used models. We only included models for which model weights are freely available and that could be run on widely available hardware, such as single-GPU compute nodes. The provided project repository (see Data Availability) contains Python scripts to run each model (one script per model) as well as shell scripts containing examples of how the Python scripts were run.

The sequence-only models AMPLIFY^57^, ESM-2^8^, and ESMC^58^ were run as masked language models. The models ISM^60^ and ISMC^60^ were also evaluated in sequence-only inference mode. (These models are trained with structural information but accept only sequence tokens at inference.) For all of these models, each target position was masked in turn and logits were extracted at the masked position. Only logits corresponding to the canonical twenty amino acids were retained. These logits were then converted into probabilities using the softmax function, *s*(*x_i_*) = exp(*x_i_*)*/* Σ*_j_* exp(*x_j_*), where *x_i_* is the logit corresponding to amino acid *i*.

The structure-aware and hybrid models required more varied inputs. SaProt^61^ was run on sequences formed by interleaving amino-acid tokens with Foldseek 3Di tokens. In masked-language-modeling (MLM) mode, the amino-acid/3Di pair at each target position was masked using the SaProt mask token. The combined amino-acid/3Di token sequence was forward passed through the model, and logits at the masked position were extracted. Only logits corresponding to the canonical twenty amino acids were retained and converted to probabilities using the softmax function.

ProstT5^62^ is a bilingual model capable of translating between amino-acid and 3Di token sequences in either direction. Here, it was used in the 3Di-to-amino-acid direction: the model was provided with Foldseek 3Di token sequences prefixed with the <fold2aa> special token. Logits at mutation position were extracted, restricted to the canonical twenty amino acids, and converted to probabilities using the softmax function.

ProteinMPNN^9^ and SolubleMPNN^63^ were run with the score.py script from the LigandMPNN repository (https://github.com/dauparas/LigandMPNN), using use_sequence=0 so that only backbone geometry features were fed as input to the model. Logits were outputted by the script and converted to probabilities for the 20 canonical amino acids using the same method as for sequence-only models.

ESM-3^59^ was evaluated in three modes: sequence-only (masked sequence tokens without structural input), structure-only (structure tokens calculated from PDB coordinates and provided without masking), and hybrid (sequence tokens masked and structure tokens supplied without masking). Structure tokens were generated using the ESM-3 structure encoder. In all three modes, logits were computed at the masked sequence positions using the respective input information, restricted to the 20 canonical amino acids, and converted to probabilities via the softmax function.

### Mutation scoring, benchmarking, and statistical analyses

To assess the quality of model predictions, we first converted all predicted amino-acid probabilities into log-odds relative to wild type. The log-odds are defined as

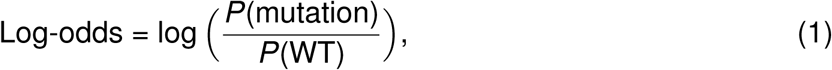

where *P*(mutation) is the predicted probability of a given mutation and *P*(WT) is the predicted probability of the wild type amino acid at the same site.

The model performance for a dataset was subsequently quantified as the Spearman rank correlation coefficient (*ρ*) between the log-odds scores and the experimental phenotype value. Correlation coefficients were computed separately for each dataset and model.

To compare experimental phenotypes with each other in the inter-experimenter and inter-phenotype analyses, we computed Spearman rank correlation coefficients separately for each dataset and phenotype pair. These correlations were computed after conducting an inner-join by mutation across all datasets for each protein. This way, only the mutations contained in both studies contributed to the correlation.

For the screening analysis in Fig. 5, we selected the ProteinGym datasets whose metadata coarse_selection_type was annotated as Activity. We retained only known experimentally assayed single mutants for which a model score could be calculated and ranked them in decreasing mutation log-odds. Within each assay, we defined nested targets as ProteinGym’s experimentally fit mutations (DMS_score_bin=1), variants at or above the 90th percentile of the experimental DMS score (top 10%), and variants at or above the 99th percentile (top 1%). Quantile thresholds were inclusive to retain tied scores.

For each screening budget *k ∈ {*8, 16, 32, 64, 128, 256*}*, we calculated target recall as the fraction of all target mutations occurring among the *k* highest-ranked candidates. We divided this recall by the expected random recall, min(*k/n*, 1), where *n* is the number of assayed mutations in that dataset, to obtain selection enrichment. We report the unweighted mean across function assays (Fig. 5B); error bars are standard errors across assays. Because an individual recovered target can yield a large recall enrichment in a large assay, this mean summarizes heterogeneous assay-level effects and should not be interpreted as a uniform fold improvement.

We then used a budget of *k* = 32 to record whether the model-ranked candidates contained at least one top-10% variant, at least one top-1% variant, or an experimental-score maximum (**Fig. 5C**). The random comparison is the probability of at least one hit under sampling without replacement from the *n* assayed variants, calculated separately for each assay. We report the mean hit probability, with standard errors across assays.

For new-to-nature datasets, raw experimental values were z-scaled per study, then normalized for comparison by placing WT at 0: pose2 percentage^53^, log_10_[(*k*_cat_*/K*_m_)_2NH_*/*(*k*_cat_*/K*_m_)_Paroxon_] with round-to-round differences^54^, trans-isomer percentage^49^, total turnover number^50^, and the maximum specificity shift across assayed substrates^51^. Model log-odds were computed using the sequence background corresponding to each directed-evolution round. For the engineering of *Pseudomonas diminuta* phosphotriesterase, we included a second analysis where log-odds at each step was computed against the wild type background (Fig. S8).

To combine datasets, each study was *z*-scored using all variants including WT, and the *z*-scores were subsequently shifted (by subtracting the *z* of the WT) such that the WT variant had *z* = 0. The *z*-scores were computed as *z_i_* = (*x_i_ − x^-^*)*/σ*, where *x_i_* is the experimentally measured phenotype value for the variant, *x^-^* is the mean over all *x_i_* for a given study, and *σ* is the corresponding standard deviation.

## RESOURCE AVAILABILITY

### Lead contact

Requests for further information and resources should be directed to and will be fulfilled by the lead contact, Claus O. Wilke.

### Materials availability

N/A

### Data and code availability

- All data and code used in the analysis for this paper is contained at the following GitHub repository: https://github.com/prwoolley/zero_shot_analysis.
- Repository stored on Zenodo: https://doi.org/10.5281/zenodo.20320202
- Associated files: https://doi.org/10.5281/zenodo.20326356

## ACKNOWLEDGMENTS

This work was supported by DTRA (HDTRA1201001) and NIH (1R01GM146093-01A1). A.D.E. is also supported by the Welch Foundation (F-1654). C.O.W. acknowledges support from the Blumberg Centennial Professorship in Molecular Evolution. A.L.F was supported in part by the Charles W. Smith Jr. Graduate Fellowship in Artificial Intelligence and Machine Learning for Biosciences.

Computational analyses were performed using the Biomedical Research Computing Facility (RRID:SCR_021979) at UT Austin, Center for Biomedical Research Support.

## AUTHOR CONTRIBUTIONS

All authors contributed to the study conception and design. Data collection, method development, analysis, and figure preparation were performed by P.R.W. and A.L.F. The first draft of the manuscript was written by P.R.W. All authors contributed to editing and revising the manuscript. All authors read and approved the final manuscript.

## DECLARATION OF GENERATIVE AI AND AI-ASSISTED TECHNOLOGIES

During the preparation of this work, the authors used an AI text editor to navigate LaTeX settings and suggest textual edits. The authors reviewed and edited the content as needed and take full responsibility for the content of the published article.

## SUPPLEMENTAL INFORMATION INDEX

Data S1. Summary of all FireProtDB datasets used in our analysis (FireProtDB_Datasets.csv)

Data S2. Summary of all ProteinGym datasets used in our analysis (ProteinGym_Datasets.csv)

## Supplementary Information

**Figure S1:**
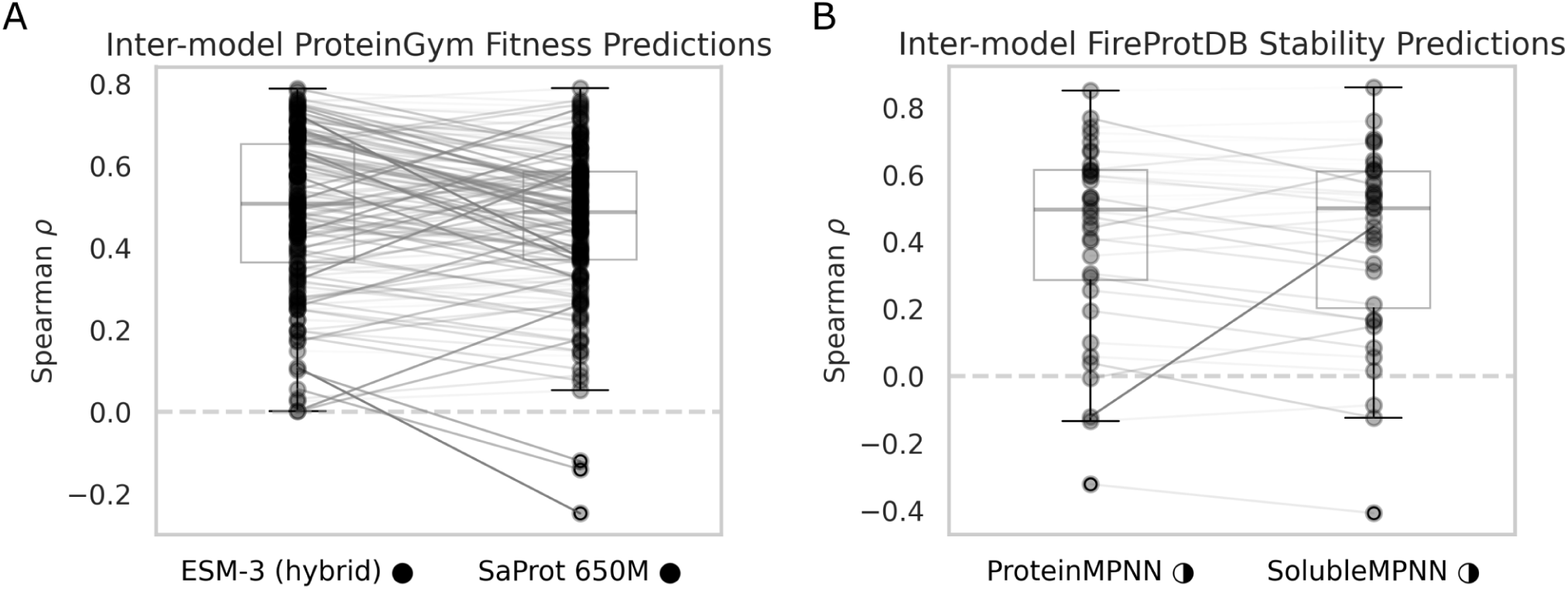
Models of comparable aggregate performance have high per-dataset variance. Zero-shot model performance (Spearman *ρ*) on the FireProtDB thermostability benchmark (A) and the ProteinGym Fitness benchmark (B). Dots represent model performance on individual datasets and lines connect the same datasets for different models. Boxplots highlight the median and interquartile range of model performance. The horizontal dashed line indicates random performance (*ρ* = 0).

**Figure S2:**
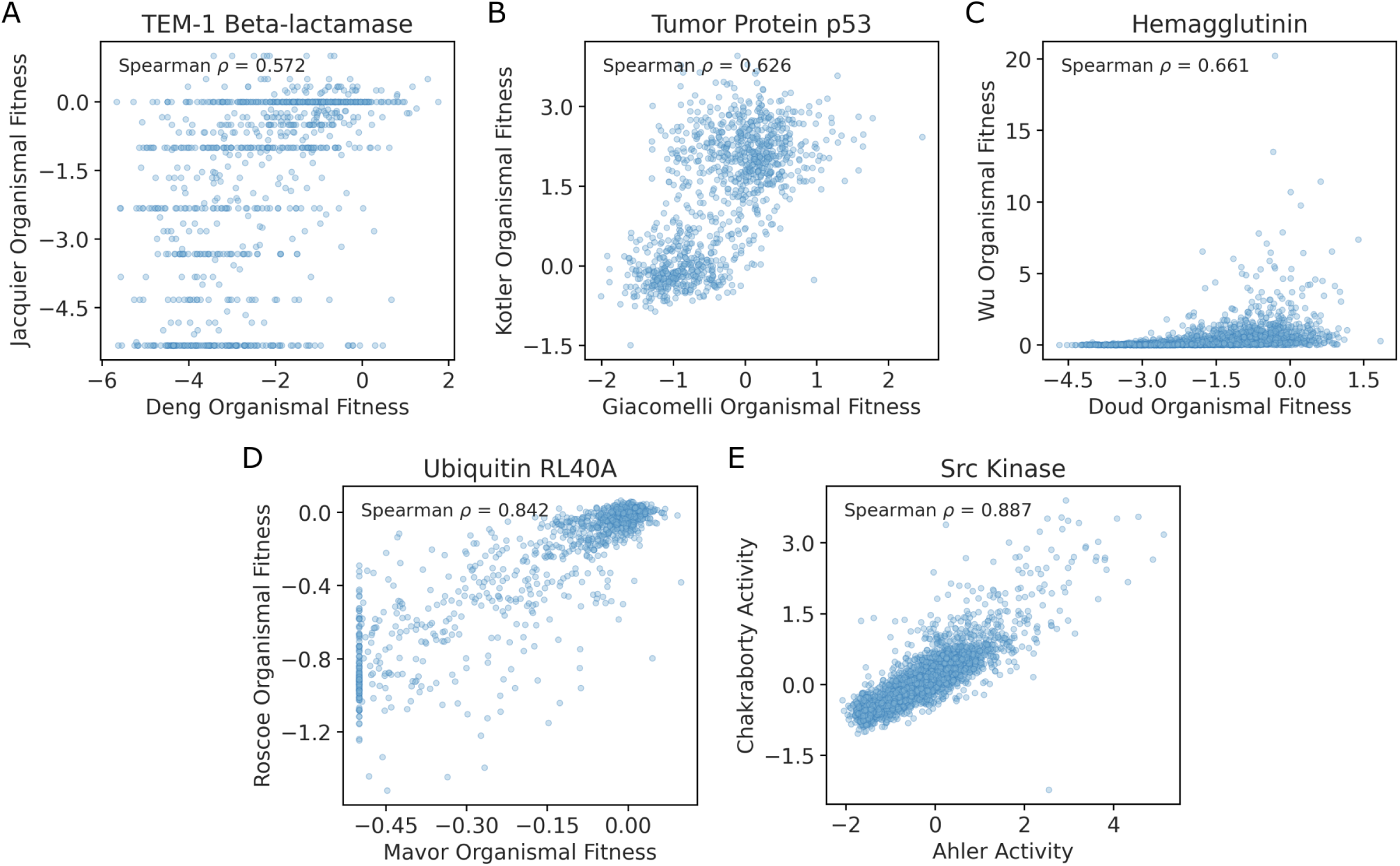
Additional inter-experimenter comparisons within ProteinGym. Scatter plots illustrate the correlation of phenotypic measurements from independent groups assaying the same protein for the same objective, ordered by ascending Spearman *ρ*. (A) TEM-1 *β*-lactamase organismal fitness. (B) Tumor Protein p53 organismal fitness. (C) Hemagglutinin organismal fitness. (D) Ubiquitin RL40A organismal fitness. (E) Src Kinase activity.

**Figure S3:**
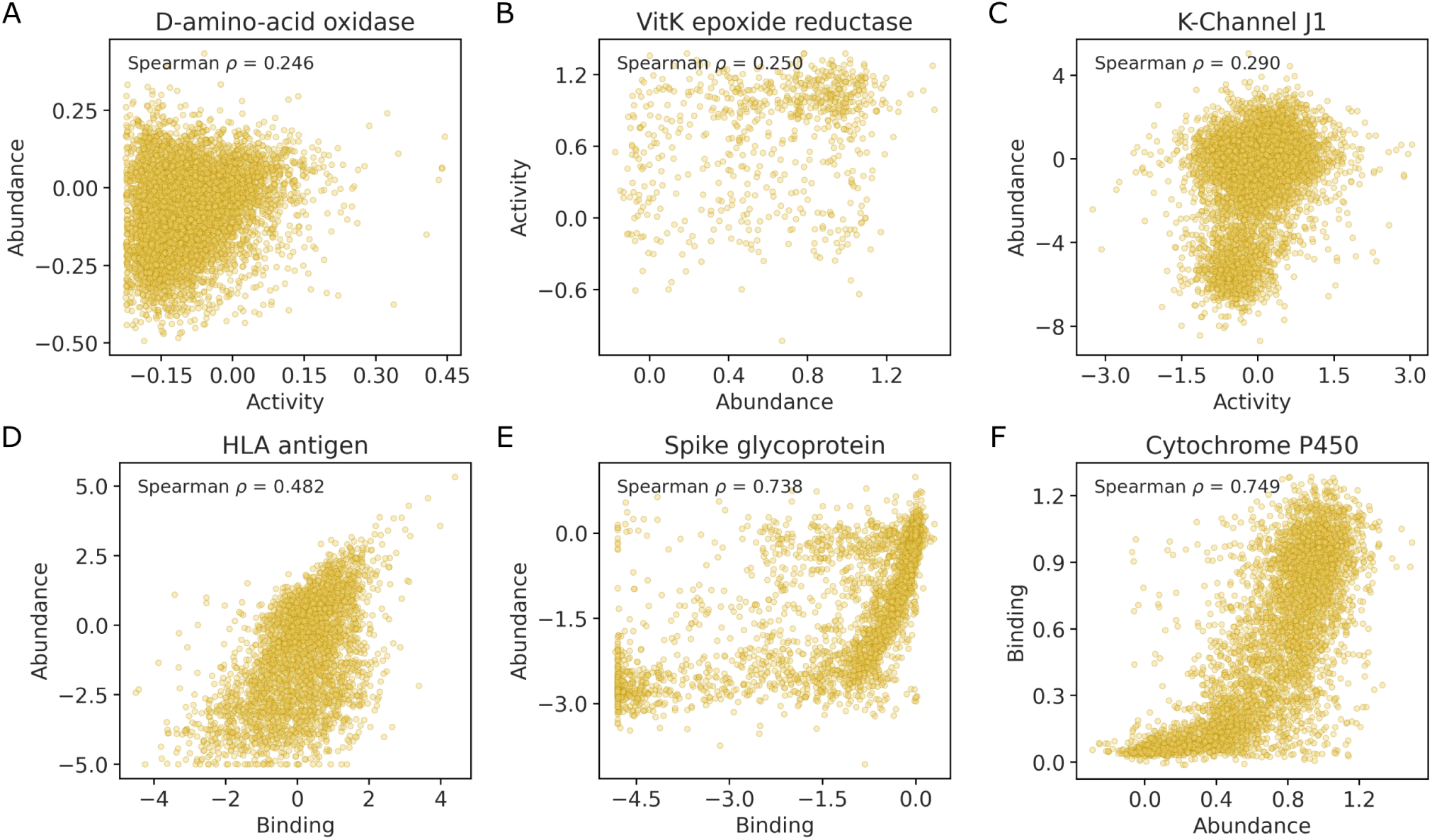
Additional inter-phenotype comparisons within ProteinGym. Scatter plots illustrate the correlation of two distinct phenotypic measurements from a single group assaying the same protein, ordered by ascending Spearman *ρ*. (A) D-amino-acid oxidase activity versus abundance. (B) VitK epoxide reductase abundance versus activity. (C) K-Channel J1 activity versus abundance. (D) HLA antigen binding versus abundance. (E) Spike glycoprotein binding versus abundance. (F) Cytochrome P450 abundance versus binding.

**Figure S4:**
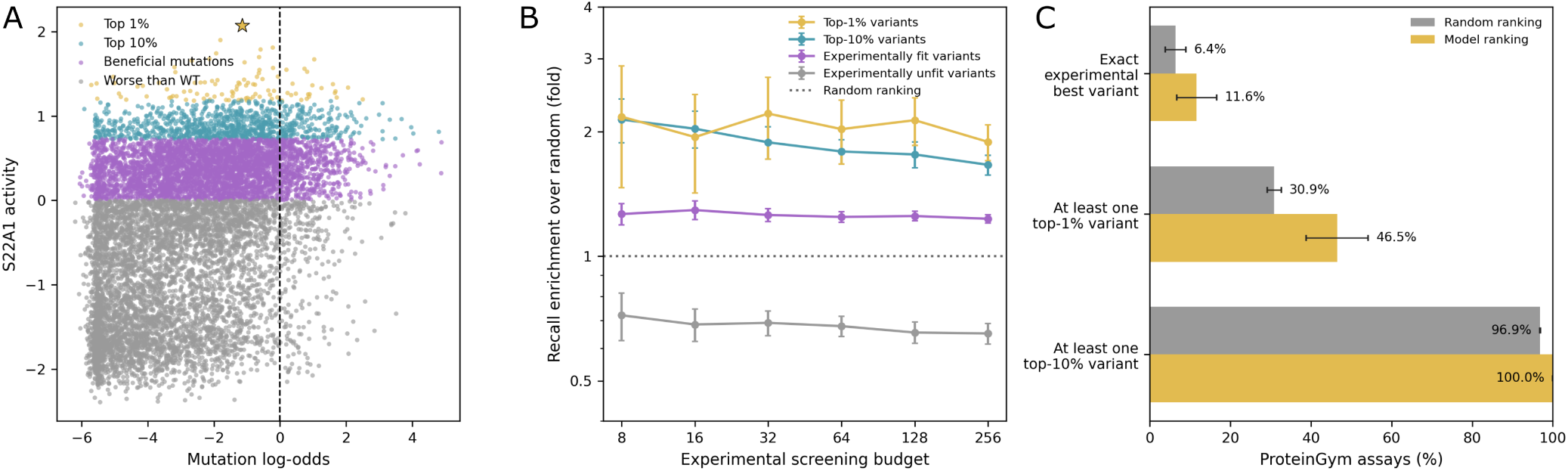
ProteinMPNN ranking enriches experimentally fit and high-scoring variants in ProteinGym Activity assays. (A) Experimental activity versus mutation log-odds for solute carrier 22A1 (S22A1)^27^. For this wild-type-centered assay, grey points are worse than wild type, purple points are remaining beneficial mutations, blue points are in the top 10% of activity values, and gold points are in the top 1%; the star marks the experimentally best variant. The vertical dashed line indicates zero log-odds. (B) Selection enrichment, defined as the ratio of model-ranked recall to the expected recall from random screening, across six base-two screening budgets. Lines show the mean across ProteinGym datasets annotated as Activity; error bars are standard errors across assays. The top-1%, top-10%, and experimentally fit targets are nested. (C) At a fixed budget of 32 known assayed mutations, the fraction of Activity assays in which random or model ranking recovers the exact experimental best variant, at least one top-1% variant, or at least one top-10% variant. Random baselines are exact hypergeometric probabilities for each assay, and error bars are standard errors across assays.

**Figure S5:**
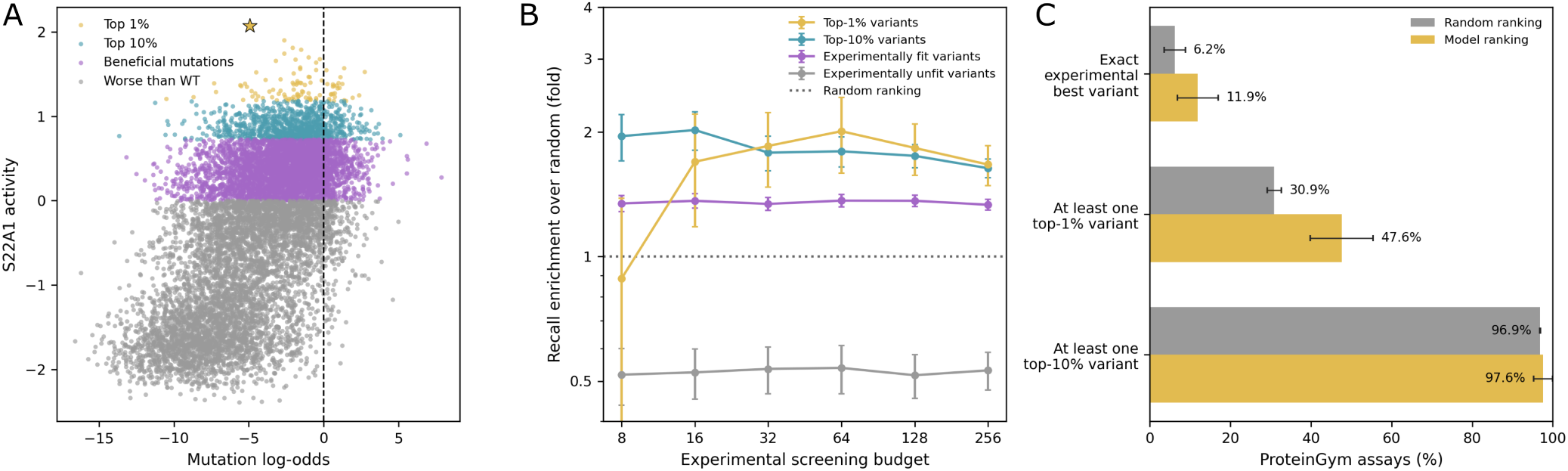
ESMC-600M ranking enriches experimentally fit and high-scoring variants in ProteinGym Activity assays. (A) Experimental activity versus mutation log-odds for solute carrier 22A1 (S22A1)^27^. For this wild-type-centered assay, grey points are worse than wild type, purple points are remaining beneficial mutations, blue points are in the top 10% of activity values, and gold points are in the top 1%; the star marks the experimentally best variant. The vertical dashed line indicates zero log-odds. (B) Selection enrichment, defined as the ratio of model-ranked recall to the expected recall from random screening, across six base-two screening budgets. Lines show the mean across ProteinGym datasets annotated as Activity; error bars are standard errors across assays. The top-1%, top-10%, and experimentally fit targets are nested. (C) At a fixed budget of 32 known assayed mutations, the fraction of Activity assays in which random or model ranking recovers the exact experimental best variant, at least one top-1% variant, or at least one top-10% variant. Random baselines are exact hypergeometric probabilities for each assay, and error bars are standard errors across assays.

**Figure S6:**
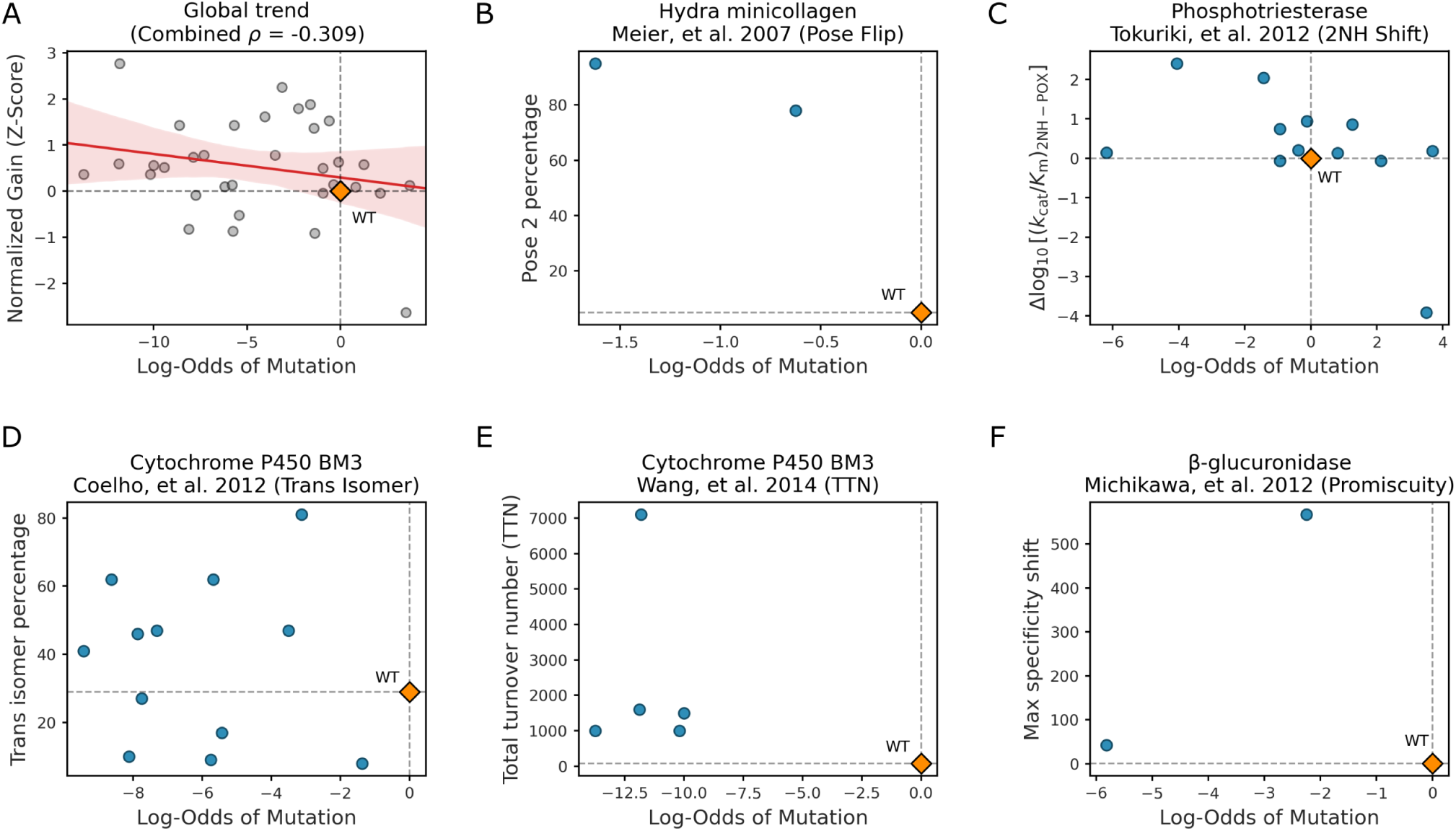
ESMC performs poorly on new-to-nature function mutations. (A) Pooled results from five examples of new-to-nature engineering campaigns. Each point represents a single mutation. Normalized gain indicates the fitness increase (or decrease) of a mutation relative to wild type, represented as a *z*-score to make results comparable across the five experimental systems. Log-odds is the model prediction for each mutation. Overall, there is a slight negative correlation between normalized gain and log-odds, i.e., mutations predicted to be better perform on average worse. (B) Hydra minicollagen engineering to favor an alternative conformational state (Meier et al. 2007).(C) Phosphotriesterase engineering to favor an alternative substrate (Tokuriki et al. 2012).(D) Cytochrome P450 engineering to optimize trans isoform percentage (Coelho et al. 2012).(E) Cytochrome P450 engineering to optimize total turnover number (TTN) (Wang et al. 2014).(F) *β*-glucuronidase engineering for promiscuity toward alternative substrates (Michikawa et al. 2012).

**Figure S7:**
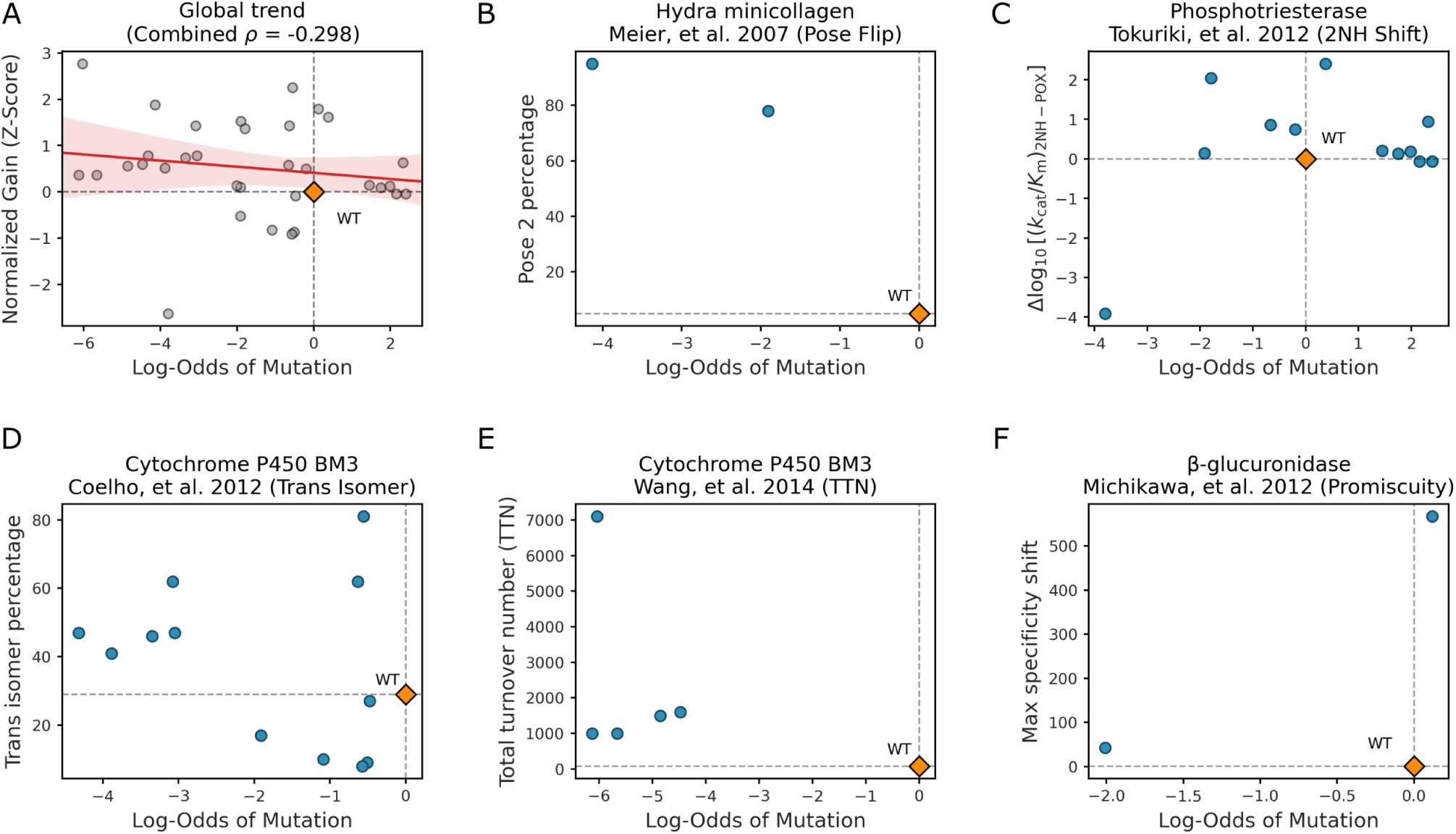
ProteinMPNN performs poorly on new-to-nature function mutations. (A) Pooled results from five examples of new-to-nature engineering campaigns. Each point represents a single mutation. Normalized gain indicates the fitness increase (or decrease) of a mutation relative to wild type, represented as a *z*-score to make results comparable across the five experimental systems. Log-odds is the model prediction for each mutation. Overall, there is a slight negative correlation between normalized gain and log-odds, i.e., mutations predicted to be better perform on average worse. (B) Hydra minicollagen engineering to favor an alternative conformational state (Meier et al. 2007).(C) Phosphotriesterase engineering to favor an alternative substrate (Tokuriki et al. 2012).(D) Cytochrome P450 engineering to optimize trans isoform percentage (Coelho et al. 2012).(E) Cytochrome P450 engineering to optimize total turnover number (TTN) (Wang et al. 2014).(F) *β*-glucuronidase engineering for promiscuity toward alternative substrates (Michikawa et al. 2012).

**Figure S8:**
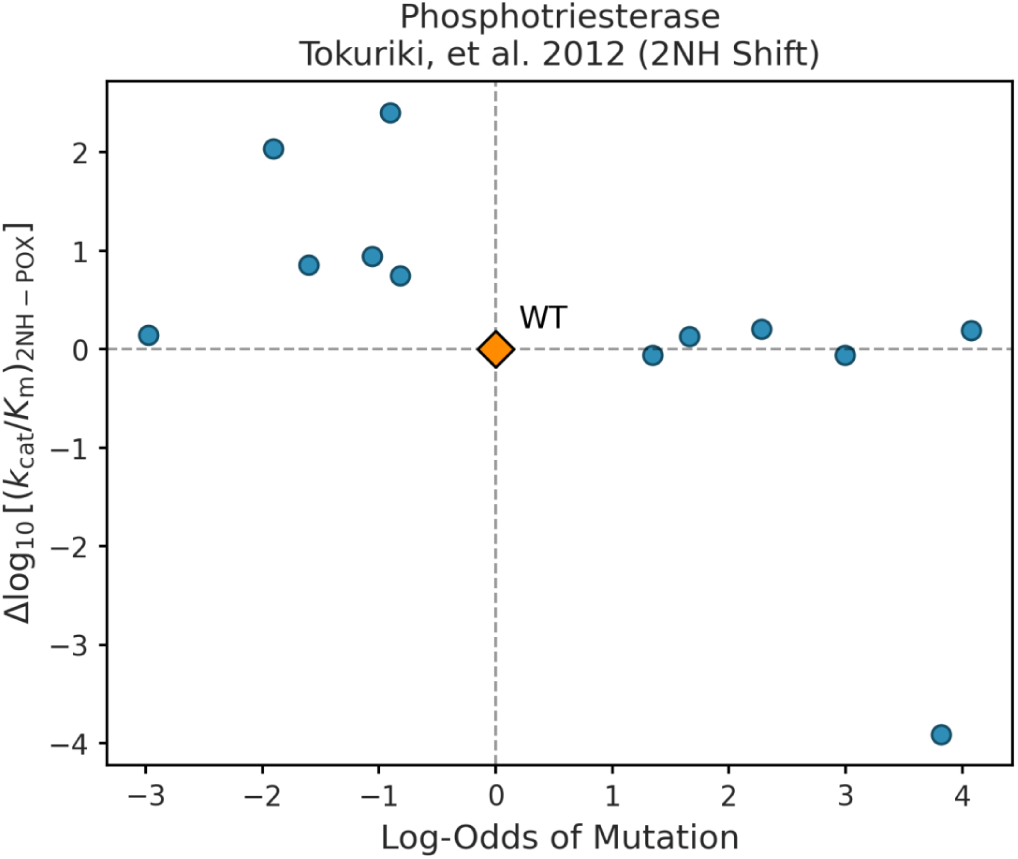
ESM3-hybrid predictions of Phosphotriesterase engineering with WT background. Phosphotriesterase engineering to favor an alternative substrate (Tokuriki et al. 2012) with mutations predicted from the WT sequence background as opposed to the previous mutation round. ESM3-hybrid mutation log-odds are plotted against the activity for the alternative substrate.

